# TRAIL orchestrates ThINKK-induced NK cell cytotoxicity against childhood acute lymphoblastic leukemia

**DOI:** 10.1101/2025.11.12.687655

**Authors:** Émilie Ollame-Omvane, Leila Ben Khemis, Paulo Cordeiro, Claire Fuchs, Alex Richard-St-Hilaire, Kathie Béland, Elie Haddad, Daniel Sinnett, Sabine Herblot, Michel Duval

## Abstract

**Background:** Therapeutic Inducers of Natural Killer cell Killing (ThINKK) represent a novel class of immunotherapy designed to enhance the graft-versus-leukemia effect of hematopoietic stem cell transplantation in pediatric patients with high-risk or relapse leukemia. Our previous work identified high expression of TRAIL as a key signature of Natural Killer (NK) cell stimulation by ThINKK. In this study, we aim to elucidate the mechanisms underlying acute lymphoblastic leukemia (ALL) killing by ThINNK-stimulated NK cells and to identify predictive sensitivity markers of this innovative approach.

**Methods:** We performed NK cell cytotoxic assays using a panel of genetically diverse ALL cell lines and patients’ samples. Gene deletion and gene enforced expression in sensitive or resistant cell lines were performed to demonstrate the role of TRAIL-receptors expression and death receptor signaling pathway in ALL cell death induced by ThINKK-stimulated NK cells. These findings were further validated through the analysis of primary patients’ samples and transcriptomic profiling of a cohort of 320 ALL patients from the CHU Sainte-Justine.

**Results:** We found that ALL sensitivity to ThINKK-stimulated NK cell killing was independent of their genetic background or their HLA expression. In addition, our data revealed the dual role of TRAIL: first, a strong NK cell activating receptor that induced rapid killing of ALL expressing TRAIL-R2, and second, a death-receptor ligand inducing ALL apoptosis following sustained engagement with its receptors. The transcriptomic analysis of ALL patients’ samples indicated that TRAIL-R2 and TRAIL-R1 are widely expressed across ALL subtypes and are not downregulated at relapse.

**Conclusion:** These findings support the use of TRAIL receptor expression as a biomarker of sensitivity to ThINKK immunotherapy and establish a mechanistic framework to guide patient stratification and therapeutic optimization.

## Introduction

Relapsed acute lymphoblastic leukemia (ALL) remains the leading cause of cancer-related mortality in children, with about 40% of patients succumbing to their disease after relapse [1,2]. Large-scale genome wide sequencing analyses have revealed the high heterogeneity and complexity of the genetic lesions of childhood ALL [3,4]. Although the identification of these alterations is crucial for guiding targeted chemotherapeutic interventions, it also underlines the challenges to find a treatment that will benefit for the majority of pediatric ALL patients, particularly for chemo-resistant and relapsed leukemia.

Allogeneic hematopoietic stem cell transplantation (HSCT) has been used for decades to cure children with relapsed or refractory acute leukemia [5,6]. HSCT success relies not only on the intensive conditioning regimen but also on the eradication of residual cancer cells by the donor-derived immune system, namely the Graft-versus-Leukemia (GvL) effect. Both innate and adaptive cytotoxic lymphocytes, i.e. Natural Killer (NK) cells and T cells respectively, contribute to the GvL effect [7,8]. Donor-derived NK cells are the first immune effectors to reconstitute from donor’s hematopoietic stem cells, mediating early GvL effect, when the leukemia burden is at its lowest after the conditioning regimen [9-11]. In addition, allogeneic NK cells do not cause graft-versus-host disease (GvHD), the primary life-threatening complication of allogeneic HSCT. Indeed, GvHD is driven by donor’s T cells recognizing histocompatibility antigens on normal cells [12]. Thus, harnessing NK cell anti-leukemia activity has emerged as a promising approach to eradicate leukemia residual cells in the first months after transplant, and thus to improve HSCT patients’ outcomes [11,13].

A balance of activating and inhibitory signals tightly regulates NK cell killing activity in order to preserve normal cells and kill transformed or virally infected cells [14]. Activating receptors, such as NKG2D, DNAM-1, and the natural cytotoxicity receptors (NCR, NKp46, NKp44, NKp30) recognize stress-induced molecules on infected or malignant cells and trigger activating signals through intracellular immunoreceptor tyrosine-based activation motifs (ITAM). Conversely, inhibitory receptors, including NKG2A/CD94 and killer immunoglobulin-like receptors (KIR), recognize self-molecules such as HLA-E and classical HLA class I molecules respectively, and trigger inhibitory signals through immunoreceptor tyrosine-based inhibition motifs (ITIMs) [15,16]. When ITAM signals overcome ITIM signals, NK cells initiate a cytotoxic response through multiple mechanisms, including the release of cytotoxic granules containing perforin and granzymes, the engagement of death receptor pathways via the surface expression of ligands (Fas ligand (FasL) and TNF-related apoptosis-inducing ligand (TRAIL)), and the secretion of pro-inflammatory cytokines such as interferon-gamma (IFN-γ) and tumor necrosis factor (TNF) [16]. Despite this killing armamentarium, ALL are often resistant to resting NK cell cytotoxicity due to the low expression of NK cell receptor ligands, and/or the high expression of HLA class I molecules [17]. Of note, the resistance of ALL to the early GvL effect mediated by NK cells is responsible for persistence of residual leukemia and subsequent relapse.

To circumvent the leukemia resistance to NK cell killing, various strategies have been proposed to both lower NK cell activation threshold and increase their killing capacity [18-20]. We demonstrated that Toll-like receptor (TLR)-9 stimulated plasmacytoid dendritic cells (PDC) are stronger inducers of NK cell cytotoxicity against ALL than myeloid dendritic cells (DC), monocyte derived DC, or cytokines (IL-15, IL-2 or IFN-α)[21-23]. However, we found that PDC numbers are low in the first months after HSCT and their functions are impaired, precluding their direct stimulation in HSCT patients[24]. We therefore designed a method to expand and differentiate CD34^+^ umbilical cord blood progenitors in PDC for adoptive transfers [21]. We named these PDC surrogates Therapeutic Inducers of Natural Killer cell Killing (ThINKK), as we showed that ThINKK-stimulated NK cells exhibit a strong *in vitro* cytotoxicity against childhood ALL and other pediatric malignancies, and express a high levels of TRAIL on their surface [21,23,22,25].

To prepare the clinical translation of these findings, we aimed to determine the sensitivity of ALL carrying diverse genetic alterations and investigate ThINKK-induced cytotoxic mechanisms. We uncovered the crucial role of the TRAIL/TRAIL-R axis in ALL killing by ThINKK-stimulated NK cells and defined patients’ selection criteria to optimize the therapeutic use of ThINKK.

## Materials & methods

### Cell lines and culture conditions

Seven cell lines derived from relapsed patients with childhood pre-B cell ALL were obtained from the American Type Culture Collection (ATCC, Manassas, VA): NALM-16, SEM, RCH-ACV, REH, 380, SUP-B15, and 697 (Table 1). Cells were maintained at 37°C in a humidified incubator with 5% CO_2_. NALM-16, RCH-ACV, REH, and 697 cells were cultured in RPMI-1640 medium (Wisent, Saint-Bruno, QC, Canada) supplemented with 10% heat-inactivated fetal bovine serum (FBS; Wisent, Canada). The 380 cell line was maintained in RPMI-1640 medium supplemented with 20% heat-inactivated FBS. SEM cells were cultured in Iscove’s Modified Dulbecco’s Medium (IMDM; Wisent, Canada) with 10% heat-inactivated FBS, while SUP-B15 cells were maintained in McCoy’s 5A medium (Wisent, Canada) with 10% heat-inactivated FBS. All cell lines were mycoplasma-free as assessed by routine tests using the PCR Mycoplasma Detection Kit (Applied Biological Materials Inc.).

**Table 1.**

| Name | Subtype | Sex | Age at diagnosis | Cytogenetic and genomic features | Relapsed ALL | HLA typing | Haplotype |
| --- | --- | --- | --- | --- | --- | --- | --- |
| NALM-16 | B ALL | F | 12 Y | Near Haploid | Yes | HLA-B<sup>Bw4+Thr80</sup>; HLA-C<sup>Asn80</sup> | Bw4 - C1 |
| SEM | B ALL | F | 5 Y | t(4;11) - KMT2A-AFF1 | Yes | HLA-B<sup>Bw4+Ile80</sup>; HLA-C<sup>Asn80</sup> | Bw4 - C1 |
| RCH-ACV | B ALL | F | 8 Y | t(1;19) - TCF3-PBX1 | Yes | HLA-B<sup>Bw4+Thr80</sup>; HLA-B<sup>Bw6+</sup>; HLA-C<sup>Lys80</sup>; HLA-C<sup>Asn80</sup> | Bw4/Bw6<br>C1 C2 |
| REH | B ALL | F | 15 Y | t(12;21) - ETV6-RUNX1 | Yes | HLA-A<sup>Bw4+</sup>; HLA-B<sup>Bw6+</sup>; HLA-C<sup>Lys80</sup> | Bw4/Bw6<br>C2 |
| 380 | B ALL | M | 15 Y | t(8;14) and t(14;18)<br>IGH-BCL2 ; MYC-IGH | Yes | HLA-B<sup>Bw4+Thr80</sup>; HLA-B<sup>Bw6+</sup>; HLA-C<sup>Lys80</sup>; HLA-C<sup>Asn80</sup> | Bw4/Bw6<br>C1 C2 |
| 697 | B ALL | M | 12 Y | t(1;19) - TCF3-PBX1 | Yes | HLA-A<sup>Bw6+</sup>; HLA-C<sup>Asn80</sup> | Bw6 - C1 |
| SUP-B15 | B ALL | M | 9 Y | t(9;22) - BCR-ABL1 | Yes | HLA-A<sup>Bw4+</sup>; HLA-C<sup>Asn80</sup> | Bw4 - C1 |
| 19-2024 | B ALL | F | 5Y | ETV6-RUNX1 like | No | ND | ND |
| 21-2014 | B ALL | M | 3 Y | ETV6-RUNX1 like | No | ND | ND |
| 03-2015 | B ALL | F | 5 Y | t(12;21) - ETV6-RUNX1 | No | ND | ND |
| 06-2014 | B ALL | M | 16 Y | IKZF1 (N159Y) | No | ND | ND |

### Gene editing and TRAIL-R2 transduction in ALL cell lines

Gene deletion of TRAIL-R2 in the REH cell line was performed using CRISPR-Cas9 technology by the gene editing platform of CHU Sainte-Justine research center. For stable transfection of 697 cell line, *TNFRSF10B* coding sequence was cloned in a lentiviral vector. After clonal expansion of the transfected 697 cells, genomic sequencing of the *TNFRSF10B* transgene revealed that all isolated clones (6/6) expressed either a truncated or a mutated version of the transgene. All identified mutations affected the cytoplasmic domain of the receptor, thereby disrupting its pro⍰apoptotic signaling function. Notably, the E10⍰697 clone expresses a truncated TRAIL⍰R2 isoform that lacks the entire cytoplasmic region of the protein, while retaining high⍰level expression of the extracellular domain at the cell surface. See Supplemental material for the detailed protocols.

### In vitro NK cell stimulation with ThINKK and functional assays

ThINKK were expanded and differentiated from purified cord blood CD34+ as previously described [21]. Peripheral blood samples were obtained from healthy volunteers after written informed consent was obtained in accordance with the Declaration of Helsinki and CHU Sainte Justine Institutional Review Board approval. NK cells were then negatively selected from human peripheral blood mononuclear cells (PBMC) using the EasySep Human NK Cell Enrichment Kit (Stemcell Technologies, Vancouver BC). The purity of the selected populations was always >90% as assessed by flow cytometry. NK cells were cultured for 18h in StemSpan SFEM II (StemCell technologies) in the presence or in the absence of freshly thawed ThINKK in a ratio of 1 ThINKK for 10 NK cells.

NK cell cytotoxicity assays were performed as previously described [23]. The percentage of specific lysis was defined as follow: Specific lysis (%) = [(absolute number of lived target cells in control without NK cells– absolute number of lived target cells in sample) / absolute number of lived target cells in control without NK cells] × 100.

NK cell degranulation against ALL cells was assessed via surface CD107a staining as previously described [26]. Unstimulated and ThINKK-stimulated NK cells were co-cultured with ALL cell lines at 1:1 E:T ratio in the presence of PE-conjugated anti-CD107a mAb (BD Biosciences, San Jose CA). Monensin (GolgiStop, BD Biosciences) was added at a final concentration of 6 µg/mL one hour after the start of the co-culture to prevent degradation of reinternalized CD107a. Cells were incubated for three additional hours, and then stained with APC-conjugated anti-CD56 and PE-Cy7-conjugated anti-CD3 (Biolegend).

ALL apoptosis was assessed using Propidium iodide (PI) and FITC-conjugated Annexin V staining. Early apoptotic cells were identified as Annexin V^+^/PI^−^, and late apoptotic cells as Annexin V^+^/PI^+^.

All flow cytometry acquisitions were performed on BD LSRFortessa cell cytometer (BD Biosciences) equipped with high throughput sampler. Data analysis was performed using FlowJo software (BD Biosciences).

### In vitro monocyte stimulation with ThINKK and functional assays

Monocytes were purified from NK cell-depleted PBMC by negative selection using magnetic beads according to manufacturer instructions (EasySep™ Human Monocyte Enrichment Kit without CD16 Depletion, StemCell Technologies). Monocyte purity was always above 80%. Purified monocytes were co-cultured for 18h with ThINKK in a ratio of 1 ThINKK for 10 monocytes. Cytotoxic assays were performed as above by co-culture of unstimulated and ThINKK-stimulated monocytes with Cell trace Violet-labelled REH cells at an effector:target ratio of 5:1 in 96-well V plates for 2h. Dead cells were stained with PI. Viable REH cells were counted by flow cytometry and the percentages of specific lysis were calculated as above.

### Surface phenotype analyses

The following conjugated antibodies were used to quantify the expression of NK cell activating ligands on ALL cell lines: TRAIL-R1 (DR4, CD261)-APC, TRAIL-R2 (DR5, CD262)-PE, CD95 (Fas/Apo-1)-FITC, MICA/B-PE-Cy7, CD155 (PVR)-PE-Dazzle, CD112 (Nectin-2)-PE-Cy7 (Biolegend), ULBP-1-APC, ULBP-2/5/6-PE (R&D Systems), HLA-ABC-PE (BD Biosciences). Dead cells were excluded using SYTOX™ Blue Dead Cell Stain (Thermofisher Scientific, Waltham, MA).

### Transcriptomic analysis of ALL cell lines

ALL cell line transcriptome were obtained by Next Generation sequencing of total RNA. The bioinformatics analysis was performed using the same pipeline as patients’ ALL samples described below.

### Multiplex analysis of protein secretion

We assessed cytokine and cytotoxic protein secretion using the LEGENDplex™ Human CD8/NK Panel V02 (BioLegend, USA), a bead-based multiplex immunoassay, according to the manufacturer’s protocol. Unstimulated and ThINKK-stimulated NK cells were co-cultured for 2 hours in the presence of target cells at a ratio 1:1 in 96 wells plates. Supernatants were collected, and stored at −20⍰°C until use for analysis.

### Transcriptomic analysis of ALL samples from our clinic

ALL samples were collected from bone marrow or peripheral blood of preB-ALL patients enrolled in the Quebec childhood ALL (QcALL), Triceps and Signature cohorts at diagnosis and relapse. Library preparation was conducted following Illumina’s TruSeq Stranded Total RNA protocol, which included targeted removal of ribosomal RNA and globin transcripts using the Ribo-Zero Gold kit. Sequencing was performed on HiSeq 4000 (paired-end, 75 bp reads) and NovaSeq 6000 (paired-end, 100 bp reads) platforms, achieving a mean depth of ∼150 million reads per sample. Bioinformatic analysis: ribosomal RNA (rRNA) were removed using bowtie2 [27]. Alignment to the hg19 (GRCh37) reference genome was performed by using the Spliced Transcripts Alignment to a Reference (STAR) aligner (v 2.5.3a) [28]. Gene count was measured with the HTSeq software [29] using the Ensembl version 75 gene annotation. Batch effect between cell lines and patients data was corrected using ComBat [30] and gene counts were normalized with DESeq2 [31].

### Statistical analyses

Statistical analyses were performed using Prism (GraphPad Software, San Diego, CA). Paired t-tests were used for the comparison of paired samples and one-way ANOVA for multiple comparisons. A value of p<0.05 (*) was considered significant with a confidence interval of 95%.

## Results

### Childhood pre-B ALL sensitivity to ThINKK-stimulated NK cells is independent of their genetic alteration and HLA class I KIR-related subgroup

Pre-B ALL is a genetically heterogeneous blood cancer with diverse genetic abnormalities. We therefore aim to extend our previous analysis of ALL killing by ThINKK-stimulated NK cells to a new series of relapsed pre-B ALL [23]. These cell lines have been selected for their diverse cytogenetic abnormalities and HLA haplotypes (Table 1). Cytotoxic assays against ALL were performed using purified blood NK cells from healthy volunteers either unstimulated of stimulated for 18h with ThINKK. NK cell cytotoxicity against 6 out of 7 ALL cell lines was significantly increased following ThINKK-induced NK cell stimulation, although with different kinetics, suggesting specific lytic pathways depending on the target cells. Short term contact (2h) with NALM-16, SEM, REH and RCH-ACV cell lines resulted in 55-80% ALL cell death at an E:T ratio of 5:1 (Fig. 1A). However, 380, SUP-B15 and 697 displayed moderate to no sensitivity to ThINKK-stimulated NK cells after 2h of co-culture. Extending the co-culture to 24 hours increased 380 and 697 lysis, although SUPB15 remained resistant to NK cell-mediated killing. Of note, unstimulated NK cells failed to induce ALL lysis even after prolonged co-culture (Fig. 1B).

**Figure 1.**
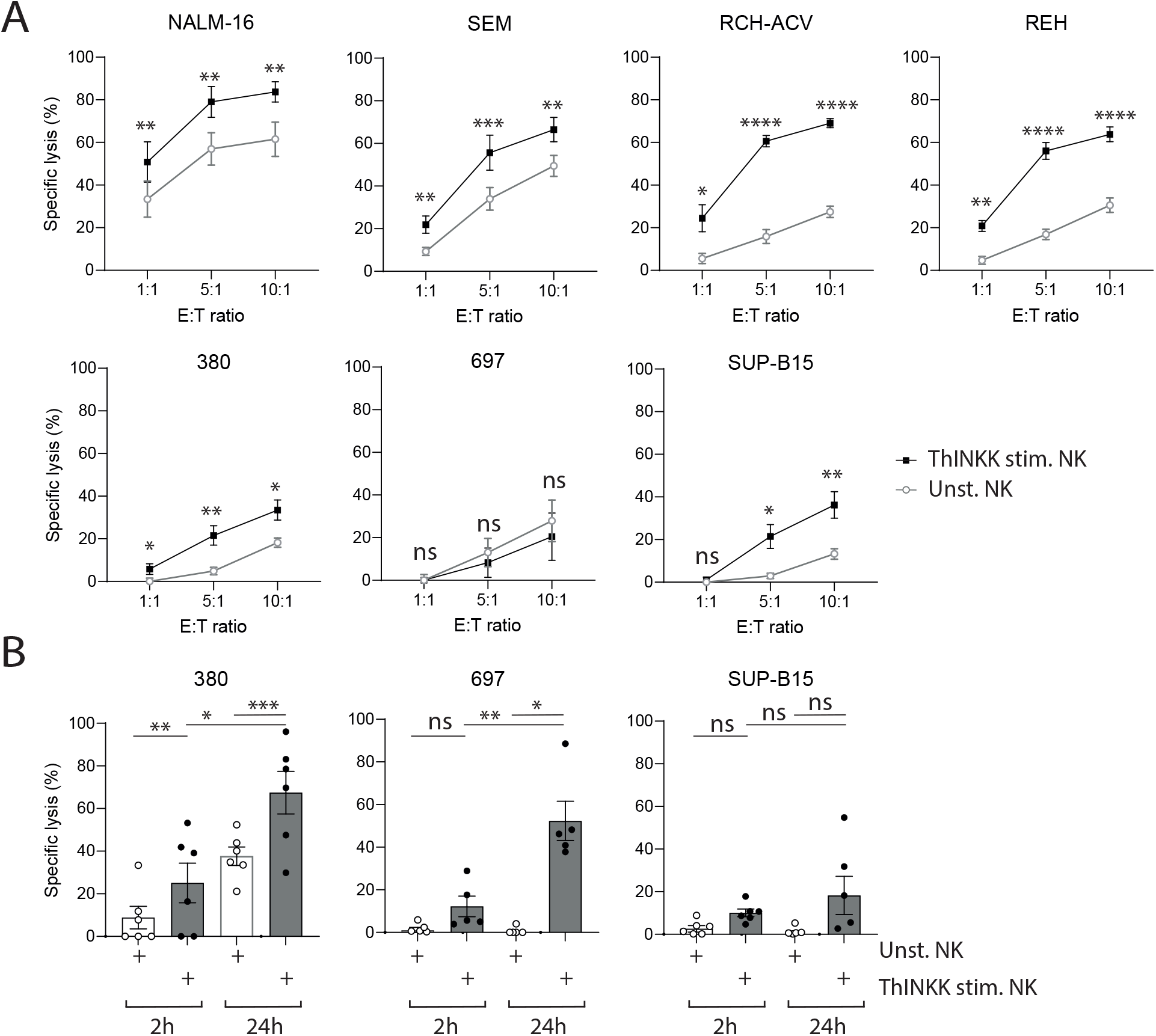
ThINKK-induced NK cell cytotoxicity against pediatric ALL killing is independent of leukemia subtype. Seven childhood ALL cell lines were selected for their diverse genetic alterations. Peripheral blood NK cells were purified from healthy volunteers and stimulated overnight with ThINKK or unstimulated. A – NK cell in vitro cytotoxic assays were performed against ALL for 2 hours. Percentages of specific lysis with SEM are represented at 3 Effector:Target (E:T ratio, 1:1, 5:1 and 10:1) (n=5 independent experiments) for the indicated cell lines. B – Specific lysis of 380, 697 and SUP-B15 were assessed after 2h (white bars) and 24h (grey bars) to evaluate late cytotoxicity of unstimulated and ThINKK-stimulated NK cells (E:T ratio of 5:1, n=5-6 independent experiments). One Way Annova test was used for multiple group comparisons **** p<0.0001, *** p<0.001, **p<0.01, *p<0.05.

In our series of ALL cell lines, the sensitivity to ThINKK-stimulated NK cell killing and the kinetics of leukemia cell death were independent of their genetic alterations or their HLA-I KIR-related subgroup (Table 1). Indeed, although RCV-ACH and 697 cell lines both harbor the same TCF3-PBX1 rearrangement, they exhibited distinct sensitivity to ThINKK-stimulated NK cells. The resistant SUPB15 cell line harbors a similar BCR-ABL rearrangement as the canonical NK cell-sensitive cell line K562. We did not observe any relation between the C1/C2 haplotypes and the sensitivity to ThINKK-induced NK cell mediated killing of ALL.

Collectively, these results indicate that although ThINKK stimulation significantly enhances NK cell cytotoxicity against most pre-B ALL, diverse killing mechanisms may be involved depending on the target. However, these mechanisms appear to be independent of the leukemia’s genetic abnormality.

### Multiple killing pathways drive ThINKK-induced NK cell cytotoxicity

Revealing the determinants of ALL sensitivity will lead to the definition of predictive biomarkers for patients who will benefit the most from ThINKK therapy. To this end, we aimed to decipher the mechanisms involved in ALL killing by ThINKK-stimulated NK cells. We investigated NK cell cytotoxic granule release and cytokine production triggered by ALL. NK cell stimulation with ThINKK significantly enhanced NK cell cytotoxic granule release against NALM-16, SEM, RCH-ACV, REH cell lines and to a lesser extent 380 cells (Fig. 2A). We further observed a correlation between the percentages of degranulation (CD107a^+^ NK cells) and the short term (2h) specific lysis of ALL (r=0.94, p=0.0013 - Fig. 2B). Accordingly, the dosage of cytotoxic molecules (Perforin, Granzymes A and B) in the supernatant of NK-ALL co-cultures revealed that ThINKK-stimulated NK cells secreted these molecules when co-cultured with target cells, but unstimulated NK cells did not. Of note, ThINKK-stimulated NK cells secreted higher levels of Granzymes A and B when co-cultured with REH cells (sensitive) as compared with co-culture with 697 cells (resistant) (Fig. 2C). We observed a similar secretion profile for the cytokines IFN-γ and TNF. While low to undetectable levels of these cytokines were observed in 2h co-cultures of unstimulated NK cells and target cells, ThINKK-stimulated NK cells secreted high amounts of IFN-γ and TNF when co-cultured with REH. Notably, 697 cells were unable to trigger the secretion of TNF by ThINKK-stimulated NK cells and induced lower production of IFN-γ compared to REH (Fig. 2D). Together, these results demonstrate that the NK cell stimulation with ThINKK enhances their short-term cytotoxicity by promoting cytotoxic granule release and by increasing the secretion of IFN-γ and TNF. However, target cells can modulate these functions, as exemplified by the lower NK cell responses triggered by 697 cells.

**Figure 2.**
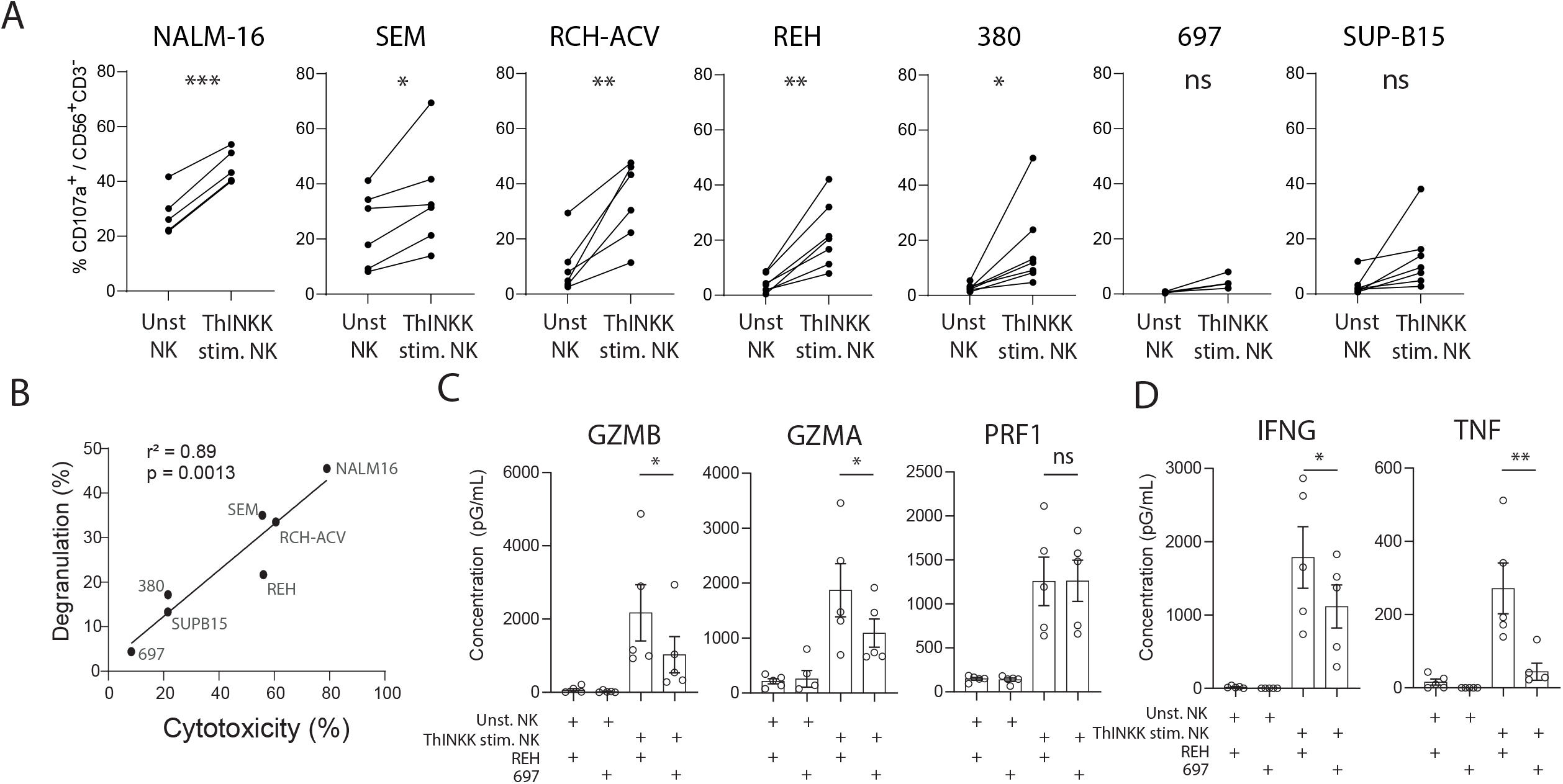
ALL sensitivity to ThINKK-induced NK cell cytotoxicity correlates with NK cell degranulation and NK cell secretion of cytotoxic molecules. A – NK cell-degranulation assays were performed against ALL cell lines using CD107a staining. Unstimulated (unst. NK) and ThINKK-stimulated NK cells (ThINKK stim. NK) were co-cultured with ALL in the presence of CD107a antibody. Percentages of CD107a NK cells are presented for 5-7 independent experiments. B – Correlation between degranulation and cytotoxicity is represented for indicated ALL cell lines. Pearson correlation test was used for statistical analysis. C-D – NK cell secreted proteins were measured in NK-ALL co-culture supernatants using a multiplex bead-based assay. REH or 697 cells were co-cultured for 2 hours with unstimulated or ThINKK-stimulated NK cells as indicated. Statistical analysis was performed using t tests to compare NK cell responses to REH and 697 cell lines.

The proportions of apoptotic ALL in co-culture with unstimulated and ThINKK-stimulated NK cells were consistent with the previous results (Fig. 3). Using Annexin V and PI staining after 2 hours of contact between NK cells and ALL, we observed that unstimulated NK cells did not trigger ALL apoptosis, except for NALM-16 and SEM, while ThINKK-stimulated NK cells increased the apoptosis of NALM16, SEM, REH and RCH-ACV. On the opposite, 380, 697 and SUPB15 cell lines did not undergo significant apoptosis within this short period of contact. Nonetheless, extending the period of contact to 24h resulted in increased apoptosis of 380 or 697. Conversely, SUPB15 did not show any increase in the proportion of Annexin V+ cells even after 24h of culture with ThINKK-stimulated NK cells.

**Figure 3.**
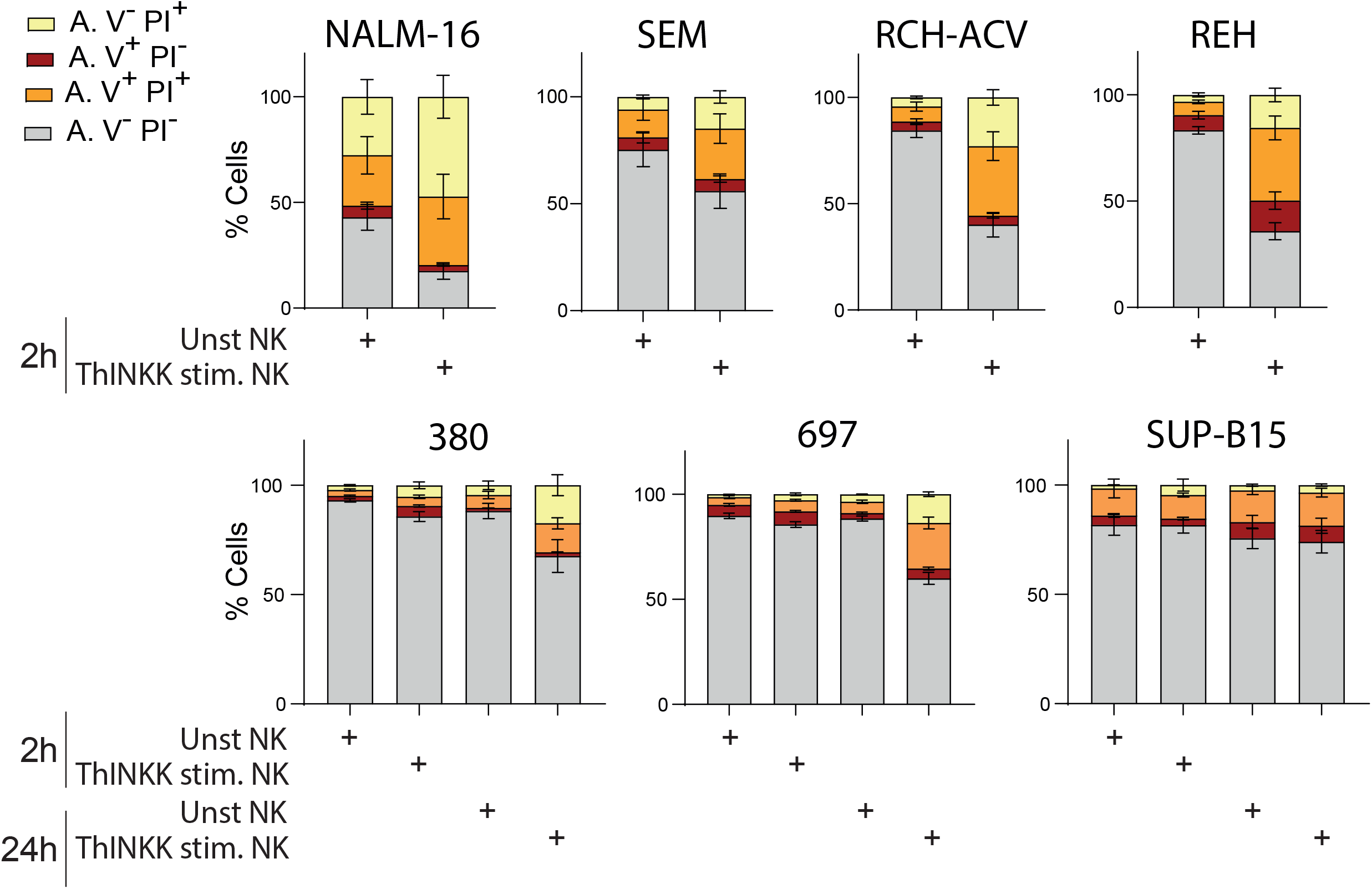
ThINKK-stimulated NK cells induce apoptosis in ALL cells with kinetics similar to cytotoxicity. ALL were stained with Annexin V and PI after 2h-or 24h-incubation with unstimulated (Unst NK) or ThINKK-stimulated NK cells (ThINKK stim. NK). Bar graphs represent the average percentages with SEM of at least 5 experiments. Non-apoptotic cells are Annexin V negative (A.V^−^ PI^−^ and A.V^−^ PI^+^) and apoptotic cells are Annexin V positive (A.V^+^PI^-^ and A.V^+^PI^+^).

Collectively, these findings demonstrate that the stimulation with ThINKK not only enhances the immediate cytotoxic functions of NK cells - evidenced by increased degranulation and secretion of IFN-γ and TNF leading to efficient apoptosis within 2 hours - but also augments delayed cytotoxic mechanisms. These delayed effects become particularly evident in target cells that exhibit resistance to short-term killing by ThINKK-stimulated NK cells.

### Pre-B ALL characteristics influencing their sensitivity to ThINKK-stimulated NK cell cytotoxicity

To identify ALL features influencing NK cell recognition, inhibition, or cytotoxicity mechanisms mediated by ThINKK-stimulated NK cells, we analyzed the surface phenotypes and the transcriptomes of pre-B ALL cell lines. We focused our analysis on the expression of class I HLA molecules, NK cell receptor ligands, and death receptors as these surface molecules directly influence target sensitivity to NK cell mediated killing. Although pre-B ALL expressed high levels of class I HLA molecules, there was no relation between the levels of expression of HLA-ABC at the surface of ALL and their sensitivity to ThINKK-stimulated NK cells (Fig. 4A-B). ALL cell lines expressed low levels of NK cell activating receptor ligands (Fig. 4A and 4C), except for NALM-16, the most sensitive cell line to NK cell killing that exhibit higher levels of PVR and ULPB-2 ligands at their surface (Fig. 4C). ALL cell lines expressed significant levels of TRAIL-R2, except 697 and SUPB15, as previously described by others for 697 (Fig. 4A and 4D)[32]. Of note, 697 expressed TRAIL-R1 while SUP-B15 did not.

**Figure 4.**
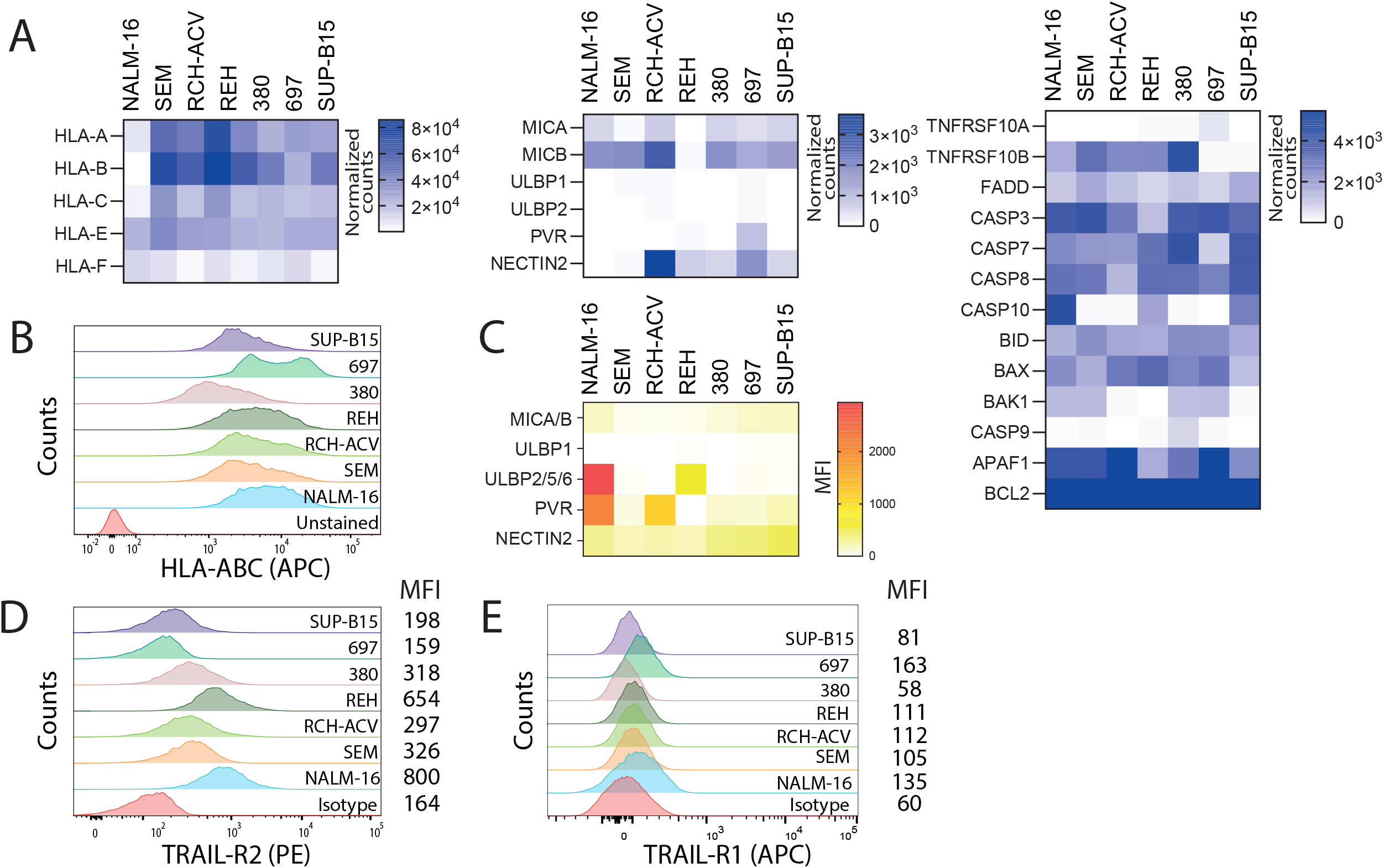
The phenotypic comparison of ALL cell lines reveals the lack of TRAIL-R2 expression in 697 and SUP-B15 as the major difference between this series of leukemia. A – Differential expression analysis of ALL cell line transcriptomes did not reveal major differences in HLA expression levels, NK cell activating receptor ligands, nor death receptor-mediated apoptosis pathway, except for TRAIL-R2 expression. Heatmaps represent normalized counts for indicated genes. B – HLA class I molecule expression was assessed by flow cytometry. Representative overlaid histograms are displayed for indicated cell lines. C – Heatmap represents the Mean Fluorescence Intensity (MFI) for indicated NK cell activating receptor ligands on ALL cell lines. D – TRAIL-R2 expression was assessed by flow cytometry. Representative overlaid histograms are displayed for indicated cell lines and isotype control.

The analysis of ALL transcriptomes did not reveal any correlation between their level of sensitivity to ThINKK-stimulated NK cells and the expression of apoptosis related genes (Fig. 4A right panel). ALL cell lines expressed genes involved in the intrinsic and extrinsic apoptosis pathways, including *FADD, CASP8* and *CASP3*. The expression of the anti-apoptotic genes such as *BCL2* or *APAF1* was elevated in ALL cell lines, but did not correlate with their sensitivity to ThINKK-induced NK cell mediated killing.

In summary, all cell lines sensitive to short-term ThINKK-stimulated NK cell killing express TRAIL-R2. Our analysis did not however find any other obvious correlations between the sensitivity to ThINKK-stimulated NK cells and ALL expression profiles.

### TRAIL-TRAIL-R2 engagement triggers multiple effector functions in ThINKK-stimulated NK cells

We previously showed that the hallmark of ThINKK-stimulated NK cells is a strong upregulation of the surface expression of TRAIL, and that TRAIL/TRAIL-R2 interaction plays a major role in ALL killing[23]. To evaluate further the role of TRAIL-R2 in NK cell-mediated cytotoxicity, we assessed the consequences of, first, the deletion of *TNFRSF10B* gene coding for TRAIL-R2 in REH, and, second, the TRAIL-R2 enforced expression on 697 cells. Flow cytometry analysis confirmed the absence of TRAIL-R2 at the surface of REH modified cell line (Fig. 5A). Importantly, the loss of *TNFRSF10B* in REH significantly reduced its sensitivity to ThINKK-stimulated NK cell-mediated lysis (Fig. 5B). Following short-term co-culture with ThINKK-stimulated NK cells, the specific lysis decreased from 38.7%±7.6 for wild type REH to 20.4%±9.3 for REH^koTRAIL-R2^ at a E:T ratio of 5:1 (p = 0.0009 paired t-test). Accordingly, the proportion of apoptotic cells decreased from 44.2%±7.6 of REH to 35.3%±4.9 of REH^koTRAIL-R2^, and NK cell degranulation decreased from 15.5%±4.1 to 8.6%±1.5 (p = 0.0037 paired t-test) (Fig. 5C-D).

**Figure 5.**
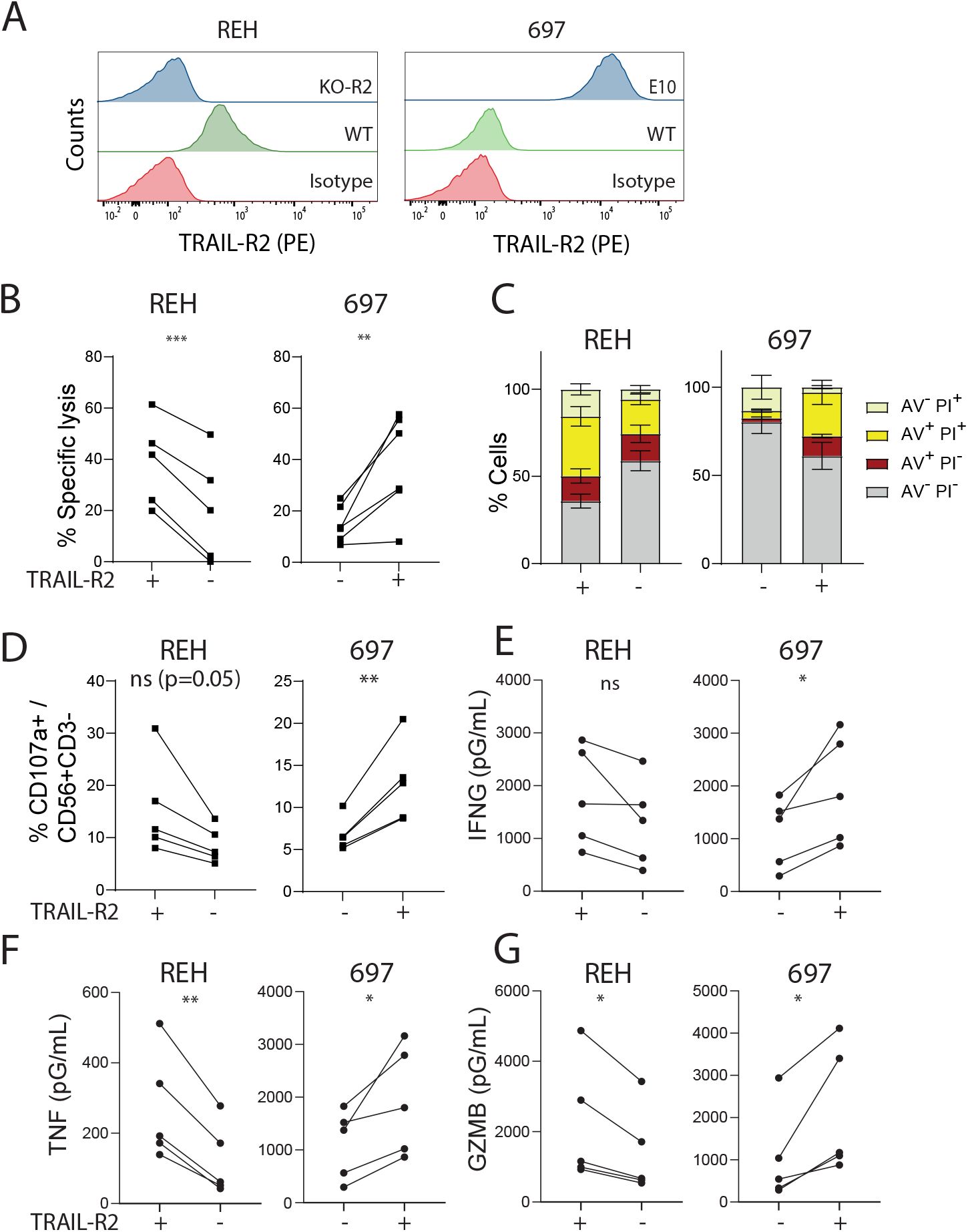
TRAIL-R2 expression governs NK cell cytotoxicity against ALL, including cytotoxic granule release, apoptosis and cytokine secretion. TRAIL-R2 expression was abrogated in REH cell line using CRISPR gene editing. Lentiviral transduction of 697 with a vector coding for TRAIL-R2 resulted in the surface expression of a truncated form of TRAIL-R2 lacking the cytoplasmic domain. ThINKK-stimulated NK cells were cultured for 2 hours with REH or 697 expressing or not TRAIL-R2 at the cell surface. A – TRAIL-R2 expression was assessed by flow cytometry on engineered ALL cell lines, REH and CRISPR-KO TRAIL-R2 REH (left panel), 697 and 697 transduced with TRAIL-R2 expressing lentivirus (clone E10-697 right panel). B – The graphs illustrate the specific lysis of REH and 697 cells expressing or lacking TRAIL-R2. C – Mean percentages of apoptotic REH and 697 cells expressing or lacking TRAIL-R2 are presented as bar graphs. D – The percentages of CD107a^+^ ThINKK-stimulated NK cells are presented in co-cultures of NK cells with REH and 697 expressing or lacking TRAIL-R2. E-G – IFN-γ, TNF and Granzyme B secretion by ThINKK-stimulated NK cells were measured using multiplex bead-based assays in supernatants of ALL-NK cell co-cultures. Statistical analysis was performed using paired t test to compare the results obtained with ALL expressing or lacking TRAIL-R2, **p<0.01, *p<0.05, ns p>0.05.

697 transduction with lentiviral vector coding for *TNFSF10RB* restored the surface expression of TRAIL-R2 (Fig. 5A), although western blot analysis and genomic sequencing of the TRAIL-R2 expressing 697 clones revealed that none of the transfected clones expressed a wild type form of this receptor. Indeed, 6 clones were sequenced and all of them exhibited either point mutations in the cytoplasmic domain or a deletion beyond the transmembrane domain (Supplemental Fig. S1). Nonetheless, despite the expression of a truncated form of TRAIL-R2, E10-697 clone exhibited an increased sensitivity to ThINKK-induced NK cell cytotoxicity as compared to untransfected 697 cells. Indeed, at a ratio E:T of 5:1, 697 specific lysis increased from 2.7%±4.6 to 32.2%±8.1 for parental and TRAIL-R2-expressing cells respectively (p = 0.003) (Fig. 5B). Conversely, the percentages of E10-697 cells apoptotic cells were increased as compared with parental 697 cells following co-culture with ThINKK-stimulated NK cells, despite the absence of the cytoplasmic death domain of TRAIL-R2 (6.3%±1.7 to 36%±7.9, p = 0.006) (Fig. 5C). The percentages of CD107a^+^ NK cells were accordingly increased in the presence of 697 expressing TRAIL-R2 as compared with parental 697 cells (6.8%±0.9 to 12.9%±2.1 p = 0.02,) (Fig. 5D). Interestingly, the production of IFN-γ, TNF and GZMB by ThINKK-stimulated NK cells was significantly lower when NK cells are exposed to TRAIL-R2 negative targets as compared to TRAIL-R2 expressing counterparts (Fig. 5E-G).

In summary, our findings establish that TRAIL⍰R2 expression on target cells is a critical determinant of ThINKK-induced NK cell cytotoxicity, driving cytotoxic granule release as well as IFN⍰γ and TNF secretion following brief effector-target interactions. Importantly, our results also reveal that the cytoplasmic death domain of TRAIL⍰R2 is dispensable for short-term ThINKK⍰induced NK cell cytotoxicity. These results reveal an unexpected, non⍰canonical function of TRAIL⍰R2, whereby its extracellular domain operates as an NK cell-activating receptor that promotes NK cell degranulation and cytolytic function independently of death domain signaling.

### Brief interaction between ALL and ThINKK-stimulated NK cells fails to activate TRAIL-R2 downstream apoptosis signaling

Given the essential role of TRAIL-R2 in controlling ALL killing by ThINKK-stimulated NK cells, we investigated whether its downstream apoptosis-signaling pathway contributes to ALL cell lysis following short and prolonged effector:target contact. We first assessed whether ALL cell lines were sensitive to soluble SuperKiller TRAIL (sTRAIL), a multimeric ligand for TRAIL receptors that has been shown to induce apoptosis of TRAIL-R2 and TRAIL-R1 expressing cancer cells [33]. Using increasing concentrations of sTRAIL and 2 incubation time points, we showed that a brief contact of 2h with sTRAIL did not induce ALL cell death in any cell line. Extending the incubation period to 24h induced NALM16, REH, 380, 697 and SEM cell death in a dose dependent manner and with variable sensitivities (Fig. 6A). Of note, SUPB15 cell line was resistant to sTRAIL induced cell death, which is consistent with its lack of TRAIL-R2 and TRAIL-R1 expression. Importantly, 24h-incubation with sTRAIL was unable to trigger RCV-ACH cell death despite their surface expression of TRAIL-R2 and their sensitivity to ThINKK-stimulated NK cell cytotoxicity. On the opposite, the lack of TRAIL-R2 expression in 697 did not impaired their sensitivity to sTRAIL after 24h of contact, confirming that TRAIL-R1 can substitute to TRAIL-R2 for apoptosis signaling, as previously published[32].

**Figure 6.**
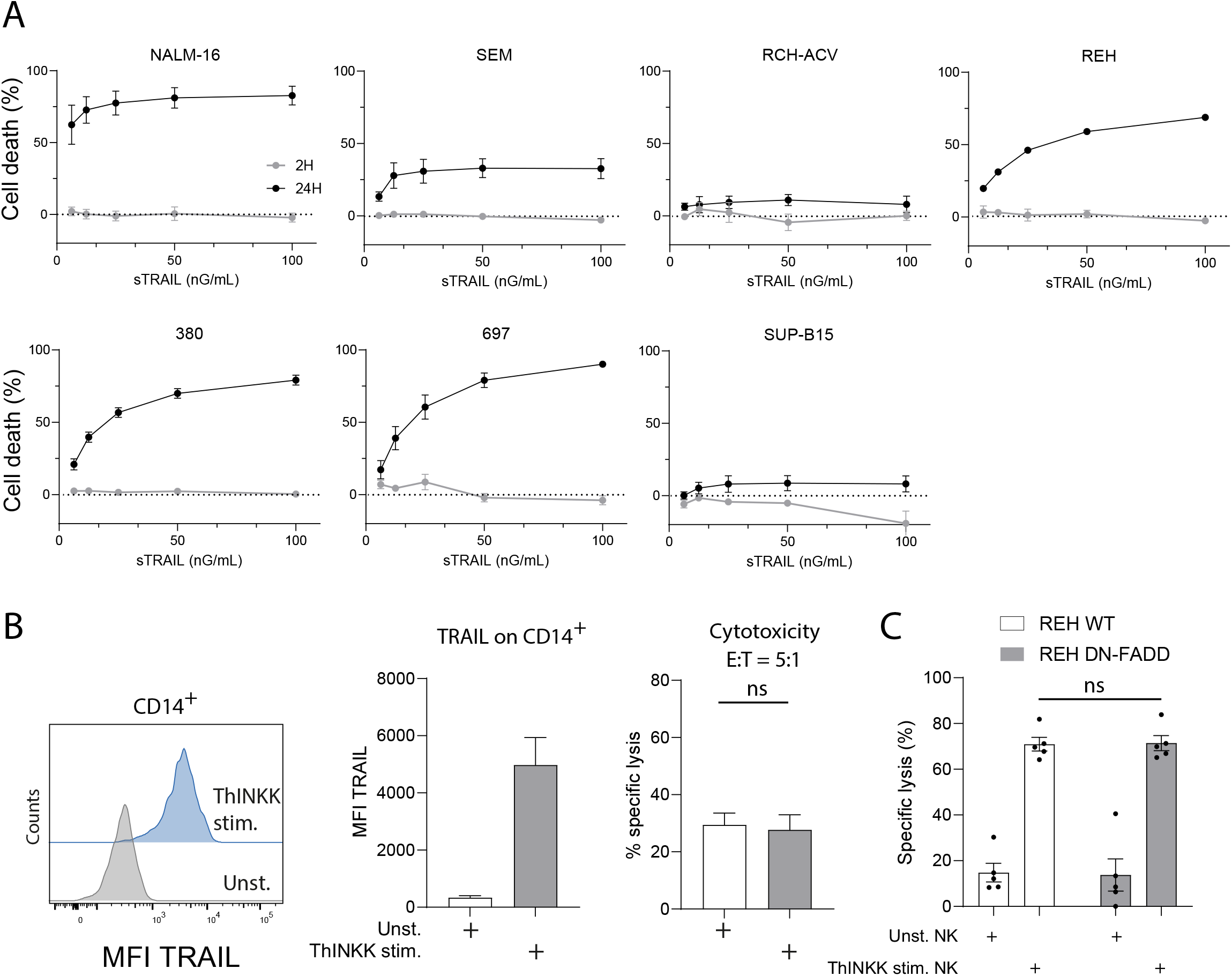
Only sustained contact between TRAIL and its receptors TRAIL-R2 and TRAIL-R1 triggers death receptor-mediated cell apoptosis. A – Indicated cell lines were cultured in the presence of increasing concentrations of SuperTRAIL (sTRAIL) for 2 hours (grey lines) or 24h (dark lines). Cell death was measured by flow cytometry after PI staining. Means of 3 independent experiments are presented with SEM. B – TRAIL expression was assessed by flow cytometry on unstimulated (unst.) and ThINKK-stimulated monocytes. Representative histograms are displayed and graph bars represents the means of 3 independent experiments. Unstimulated and ThINKK-stimulated monocytes were co-cultured for 2 hours with REH and specific lysis was assessed by flow cytometry. Bar graph represents the means of specific lysis for 3 independent experiments. C – REH cells were transduced with a lentiviral vector coding for DN-FADD. Unstimulated and ThINKK-stimulated NK cell cytotoxicity against REH wild type (WT) and expressing DN-FADD was assessed by flow cytometry. Bar graphs represent the means of 5 independent experiments with SEM. Paired t test was used for statistical comparison of samples (ns p>0.05).

Since soluble and membrane forms of TRAIL may activate distinct apoptosis signaling pathways, we took advantage of the upregulation of TRAIL surface expression on monocytes following co-culture with ThINKK to assess whether brief contact with membrane-bound TRAIL can induce ALL killing independently of cytotoxic granule release. We observed no difference in REH specific lysis when cultured for 2h with unstimulated or ThINKK-stimulated TRAIL^+^ monocytes (Fig. 6B). This result indicates that brief exposure to membrane-bound TRAIL does not induce ALL apoptosis, similarly to the effect observed with soluble TRAIL.

We next investigated the consequences of decoupling TRAIL-R2 and TRAIL-R1 from apoptosis signaling pathway by transducing REH cells with a lentiviral vector coding for a dominant negative (DN) form of FADD[34]. DN-FADD lacked the death domain preventing the activation of Caspase 8 and the downstream apoptosis cascade, as assessed by the absence of cell death induced by sTRAIL in REH expressing DN-FADD (Supplemental Fig. S2). Importantly, we observed that parental and DN-FADD-expressing REH cells exhibited the same sensitivity to short-term ThINKK-stimulated NK cell cytotoxicity (Fig. 6C).

Collectively, these results demonstrate that: (1) the death receptor-mediated apoptosis pathway requires sustained receptor-ligand interaction. (2) An intact DR-downstream signaling is not required for short-term ALL killing by ThINKK-stimulated NK cells, as exemplified by the sensitivity of RCH-ACV, REH-DN FADD and E10-697 clone expressing truncated TRAIL-R2 to ThINKK-stimulated NK cell cytotoxicity after 2h of contact. (3) In the absence of TRAIL-R2, sTRAIL binding to TRAIL-R1 mediates ALL cell death although with a delayed kinetics, such as in 697 cells. (4) The lack of both TRAIL-R1 and TRAIL-R2 expression is associated with ALL resistance to ThINKK-stimulated NK cells, such as in SUPB15 cell line.

### Potential biomarkers of patient’s leukemia sensitivity to ThINKK-induced NK cell cytotoxicity

We next took advantage of patient-derived samples from our clinic to confirm the previous results, specifically the role of TRAIL-R2 as an NK cell activating ligand, and the expression of TRAIL-R2 and TRAIL-R1 as biomarkers of sensitivity to be used as exclusion criteria in future clinical trials. We first selected four patient-derived samples according to their *TNFSR10B* gene expression levels in our transcriptomic dataset. Patients’ blasts were expanded in immune-deficient mice (patient-derived xenograft, PDX) by our in-house platform of humanized mouse models (Fig. 7A). TRAIL-R2 surface expression by these PDX was assessed by flow cytometry (Fig. 7B). Cytotoxic assays with unstimulated and ThINKK-stimulated NK cells against freshly thawed PDX samples demonstrated a good correlation between surface levels of TRAIL-R2 and PDX sensitivity to ThINKK-induced NK cell cytotoxicity (Fig. 7C-D). The limited in vitro survival of PDX cells precluded long*⍰*term cytotoxicity assays, thereby preventing assessment of specific leukemia cell lysis following extended interaction with ThINKK⍰stimulated NK cells.

**Figure 7.**
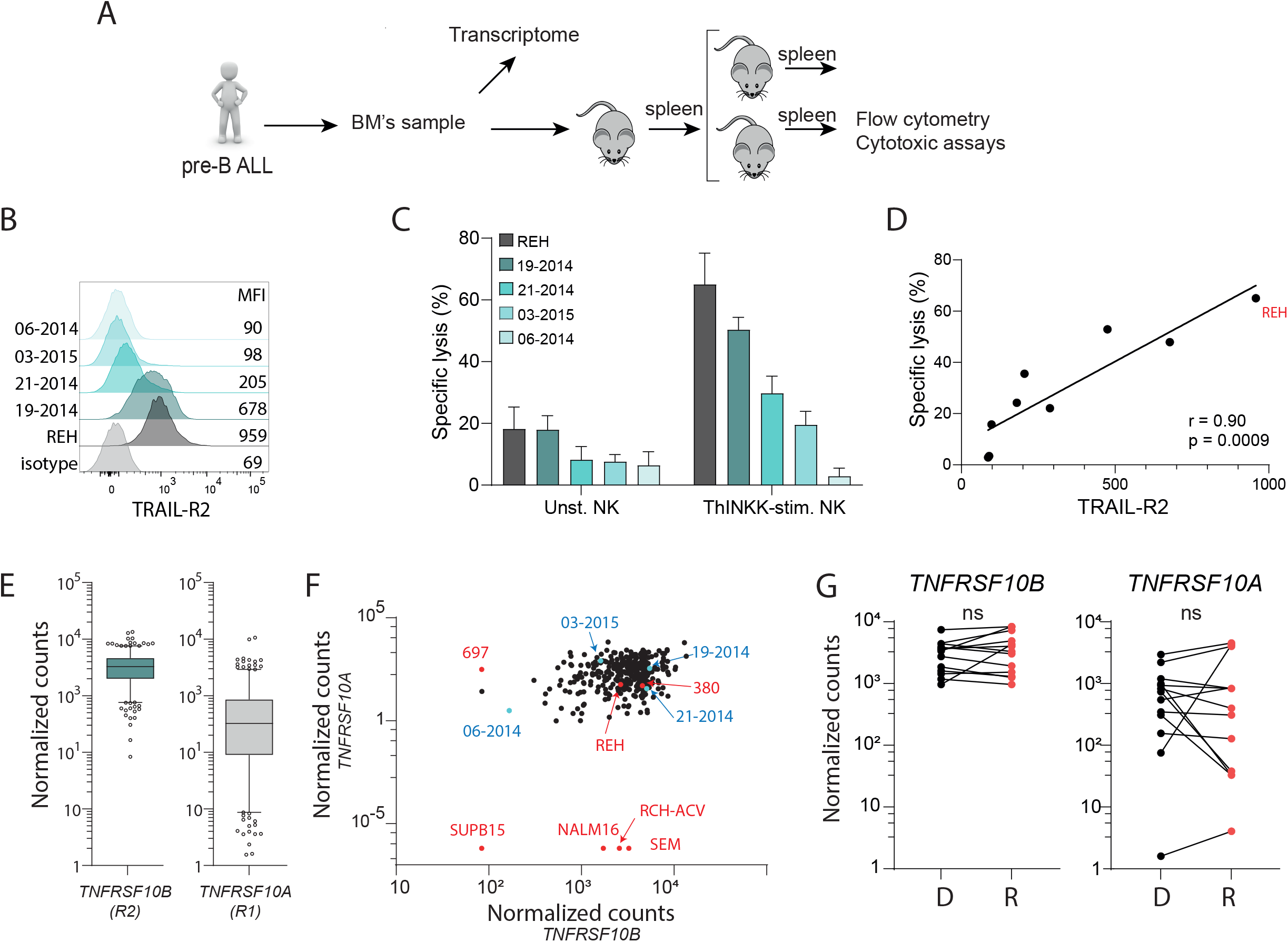
Predictive biomarkers of sensitivity to ThINKK-induced NK cell cytotoxicity. A – Schematic diagram of patient-derived xenografts (PDX) generation by *in vivo* expansion of human ALL samples in immunodeficient mice. B – The surface expression of TRAIL-R2 on PDX and REH was assessed by flow cytometry, representative histograms with Mean Fluoresence Intensity (MFI) are presented. C – Cytotoxic assays against PDX and REH (as a control) were performed with unstimulated or ThINKK-stimulated NK cells. Effector:Target ratio was 5:1 and incubation time was 2h. Means of specific lysis are presented with SEM (n=3 independent experiments). D – The correlation between the expressions of TRAIL-R2 and specific lysis is presented. E – The mRNA levels of TRAIL-R2 and TRAIL-R1 in ALL patients’ samples are presented by box and whiskers plots were the boxes represent 50% of the data where the median is presented as a line, the whiskers represent the minimum and the maximum data points and the dots represent the outliers. F – Dot plot of TRAIL-R1 (*TNFRS10A*) and TRAIL-R2 (*TNFRS10B*) mRNA levels for patients’ samples and cell lines is presented (red dots for cell lines, blue dots for PDX and black dots for all other patients’ samples). G – The mRNA levels of TRAIL-R2 and TRAIL-R1 are presented for patients’ pair of samples collected at diagnosis and relapse. Statistical analysis was performed using paired t tests (ns: non-significant).

We next investigated the expression levels of *TNFRSF10B* and *TNFRSF10A* genes (coding for TRAIL-R2 and TRAIL-R1, respectively) in a cohort of 320 pediatric pre-B ALL samples from the Division of Hematology-Oncology, CHU Sainte-Justine. Primary ALL samples (bone marrow at diagnosis or relapse) were sequenced in our institution using next generation sequencing. We integrated patients’ transcriptomic data with our dataset from ALL cell lines, and the expressions of *TNFRSF10A* and *TNFRSF10B* were analyzed across the cohort. We observed that most patients’ ALL expressed high level of *TNFRSF10B*, with a relatively narrow distribution. In contrast, *TNFRSF10A* expression showed greater variability among patient samples (Fig. 7E). Notably, none of the ALL cases displayed low expression of both *TNFRSF10B* and *TNFRSF10A*, a pattern uniquely observed in the SUPB15 cell line (Fig. 7F). Although most of our samples were collected at diagnosis, our dataset included 13 matched pairs of ALL samples obtained from the same patients at diagnosis and at relapse. The comparison of *TNFRSF10B* and *TNFRSF10A* expression levels at these two time points revealed no significant differences (Fig. 7G). These results suggest that relapsed ALL should exhibit the same sensitivity to ThINKK-induced NK cell mediated cytotoxicity as ALL at diagnosis.

Together, our findings using primary patient samples strongly support the use of TRAIL-R1 and TRAIL-R2 expression as a biomarker of sensitivity for ThINKK adoptive therapy, and in addition, reveal that most pediatric patients with ALL should benefit from this innovative therapy.

## Discussion

This preclinical study uncovers the dual role of TRAIL in childhood preB ALL killing by ThINKK-stimulated NK cells and underscores the broad therapeutic potential of ThINKK immunotherapy. Indeed, we demonstrate that TRAIL-R2 is a potent activating ligand for NK cells expressing high levels of TRAIL such as ThINKK-stimulated NK cells leading to rapid ALL killing independently of death receptor-mediated apoptosis. Additionally, the sustained engagement of TRAIL with their receptors TRAIL-R1 or TRAIL-R2 triggers ALL apoptosis through the death receptor-signaling pathway. We further reveal that these lysis mechanisms are independent of leukemia genetic alterations and that the vast majority of patients’ ALL samples express both TRAIL-R2 and TRAIL-R1.

Leukemia relapse remains a therapeutic challenge for children with high-risk acute leukemia [2,4]. Recent advances in targeted immunotherapies such as monoclonal antibodies or Chimeric Antigen Receptor (CAR)-T cells or NK cells have proven their efficacy, but also their limits. The major limits of these approaches are the immune escape by the down regulation of the target, the narrow clinical indications limited by the expression of the targeted antigen, and their toxicity related to on-target off-tumor effect [35-38]. Moreover, most of these therapies have been developed for adult hematological malignancies, causing significant challenges when translated for the treatment of young children, such as cell collection for CAR therapies or appropriate drug dosing [39]. To circumvent these challenges and limitations, we developed an innovative post-HSCT immunotherapy to improve the pediatric patients’ outcome without restriction to leukemia subtypes. Harnessing the GvL effect of HSCT offers the unique opportunity to build on the anti-leukemia activity of donor-derived immune effectors. Notably, donor-derived NK cells play a pivotal role during the early post-transplant period—when the leukemia burden is low—offering a critical window to maximize therapeutic efficacy.

Since ALL are deemed to be resistant to NK cell cytotoxicity, we engineered PDC surrogates named ThINKK, and demonstrated that ThINKK-stimulated NK cells exhibit a unique killer activity against childhood ALL and other pediatric cancers [25,22,23]. Here we extended these findings by showing that a broad variety of ALL is sensitive to ThINKK-induced NK cell killing independently of their genetic alteration or their HLA haplotype. Indeed, we identified the presence of TRAIL-R1 or TRAIL-R2 as a biomarker for ALL sensitivity and determined that their expression is not correlated with specific preB ALL subtypes. Furthermore, leukemia relapse is not associated with the downregulation of TRAIL-R1 or TRAIL-R2 [32]. We also confirm in our cohort of 320 childhood ALL patients that most of them express TRAIL-R2, and almost all express either TRAIL-R2 or TRAIL-R1.

In the absence of TRAIL-R2, TRAIL-R1 can substitute TRAIL-R2 to induce ALL apoptosis through the DR-signaling pathway, albeit with a delayed kinetic. In this context, elevated expression of apoptosis-inhibitory proteins such as BCL2, BCLXL and XIAPs, might impair TRAIL-R mediated cell death [40]. Given that pro-apoptotic agents, such as IAP or BCL2 inhibitors, are already under investigations for the treatment of childhood ALL, ongoing studies will assess their potential therapeutic benefit when combined with ThINKK-stimulated NK cells [41,42].

Our results establish the dual role of TRAIL-TRAIL-R engagement in ThINKK-induced NK cell ALL killing. We previously found that a very high expression of membrane-bound TRAIL is the hallmark of ThINKK-stimulated NK cells [22,21,23]. Furthermore, we found that TRAIL-TRAIL-R interaction plays a major role in NK cell cytotoxicity against ALL and other pediatric cancers such as neuroblastoma [22,21,23]. Using a panel of ALL cell lines and PDX we demonstrated here that, TRAIL not only induces ALL apoptosis upon sustained interaction with its receptor TRAIL-R2 and TRAIL-R1, but also is a potent activating NK cell receptor triggering a rapid cytotoxic granule release and cytokine secretion upon interaction with TRAIL-R2, independently of death-receptor downstream apoptosis signaling pathway. The role of TRAIL as an NK cell activating receptor has already been evoked in the context of viral infections. Indeed, Hofle et al. demonstrated that TRAIL contributes to NK cell anti-HIV activity independently of death receptor-mediated apoptosis, and that TRAIL engagement with TRAIL-R induces NK cell degranulation and IFN-γ secretion against HIV-infected T cells [43]. Interestingly, the authors found that decoy TRAIL receptor 1 (DcR1, TRAIL-R3) also induces NK cell activation and degranulation, despite its lack of downstream apoptosis signaling, adding a new function for this TRAIL regulatory receptor. Similarly, Vlachava et al. showed that the HCMV-derived glycoprotein gpUL4 inhibits NK cell activation and degranulation through its interaction with TRAIL at the surface of NK cells [44]. Although the downstream signaling pathway of TRAIL in NK cells remains to be fully characterized, these studies underscore the essential role of TRAIL as an NK cell activating receptor. In the context of childhood ALL, we demonstrate that TRAIL-R2 triggers this activating signal but not TRAIL-R1. Importantly, the very high expression of TRAIL on all NK cell subsets following ThINKK stimulation allows for increased NK cell responsiveness to this potent activating signal.

Although CAR T cell therapy initially demonstrated remarkable efficacy in treating pre-B ALL and lymphomas, increasing reports of relapse and resistance have emerged, resulting in a long-term survival rate of approximately 50% in children [45]. One identified mechanism of resistance involves impaired death receptor-mediated apoptosis in ALL [46,47]. Indeed, Singh et al. demonstrated that ALL cells with impaired TRAIL-R-mediated apoptosis signaling are resistant to CAR T cell cytotoxicity, and the prolonged antigen exposure induces CAR T progressive impairment of their cytotoxic potential [48]. These results underscore the requirement of both target persistence on ALL (i.e. CD19) and functional death receptor-induced apoptosis pathway for CAR-T cell-mediated killing. On the opposite, we showed here that, ALL with impaired death receptor apoptosis signaling pathway are still sensitive to ThINKK-induced NK cell killing. Since about 50% of childhood ALL have been shown to be resistant to death receptor-mediated apoptosis [48], ThINKK-stimulated NK cells should have a therapeutic advantage over CAR T, especially for patients with ALL expressing high levels of anti-apoptotic proteins.

In conclusion, our study highlights the multifunctional role for TRAIL in orchestrating NK cell effector functions and its inference in NK cell cytotoxicity against ALL. ThINKK-induced NK cell stimulation unleashes these multiple lytic functions by inducing TRAIL upregulation on all NK cell subsets. These insights not only deepen our understanding of NK cell biology in the context of ALL but also inform patient selection criteria for pediatric trials. Ultimately, they lay the groundwork for optimizing this first-in-class immunotherapeutic approach.

## Footnotes

### Declaration of interests

The authors report no conflict of interest

### Funding

This study was supported by funding from the Canadian Institutes of Health Research (CIHR #165997). EOO received a studentship from the Cole Foundation.

### Authors’ contribution

The contributions of each author are as follows: Concept and experimental plans: SH, MD, EH and DS. Experimental procedures: EOO, PC, LBK, CF, ARSH and KB. Data analysis: SH, EOO, CF, ARSH. Manuscript writing: EOO, SH, MD. Guarantors of the work: SH and MD

### Consent for publication

All authors reviewed, edited, and approved the manuscript.

### Ethics approval and consent to participate

This study was approved by the CHU Sainte-Justine Institutional Review Board (approval number: MP-21 2019-2026). All participants provided written informed consent.

### Availability of data and material

Data and material are available upon reasonable request. All data relevant to the study are included in the article or uploaded as supplementary information.

## Acknowledgements

The authors gratefully acknowledge all heathy volunteers and ALL patients for their participation to this study. We also thank all technology platforms of the Azrieli CHU Sainte-Justine Research Center that have been involved in this study, including the Gene Editing Platform for the manufacture of the CRISPR-Cas9 cell lines, the flow cytometry platform for its valuable support with flow analysis, the Next Generation sequencing platform for patients’ dataset collection. Charles Bruneau Foundation generously supports all these platforms.

## Notes

### Competing Interest Statement

The authors have declared no competing interest.

### Summary of Updates

In-depht characterization of TRAIL-R2-transfected 697 clone Figure 5 revised

