## Supplemental Fig. S1 for "TRAIL orchestrates ThINKK-induced NK cell cytotoxicity against childhood acute lymphoblastic leukemia"

### Supplemental Figure S1

A

| CLONE ID | SEQUENCING | PROTEIN |
|---|---|---|
| E10 | Stop codon 234 | Met 1 - Lys 233 |
| G10 | Full length<br>2 mutations | Pro (281) -> Leu<br>Glu (367) -> Lys |
| F5 | Stop codon 349 | Met 1 - Leu 348 |
| G4 | Stop codon 349 | Met 1 - Leu 348 |
| C1 | Full length<br>2 mutations | Ala (376) -> Ala<br>Trp (389) -> Arg |
| F7 | Stop codon 237 | Met 1 - Pro 236 |

B

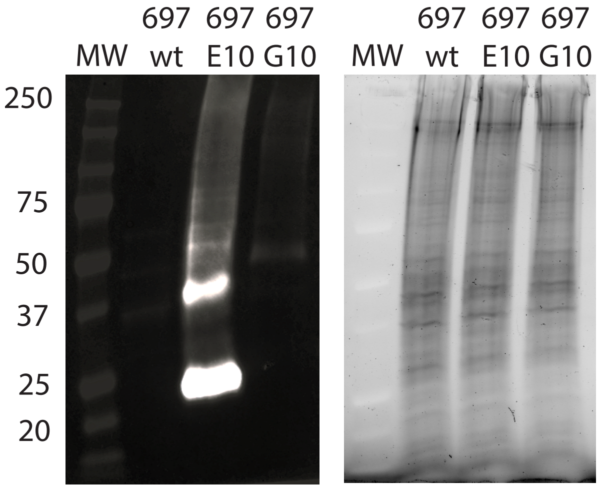


A - Sequencing of TRAIL-R2 expressing 697 clones. Six out of 6 clones of transfected 697 exhibit mutation or deletion in the cytoplasmic region of TRAIL-R2 transgene.

B - *TRAIL-R2 Western blot*. Total extracts were loaded on a 10% acrylamide mini-protean TGX stain free gel (Biorad cat 4568034), after electrophoresis and proteins transfer, anti-human TRAIL-R2 (R&D System, MAB6313), followed by goat anti-mouse secondary (Abcam, Cat ab6789) antibodies were used to reveal TRAIL-R2 proteins in 697 wild type, E10-697 and G10-697 clones.
