## Supplemental Fig. S2 for "TRAIL orchestrates ThINKK-induced NK cell cytotoxicity against childhood acute lymphoblastic leukemia"

### Supplemental Figure S2

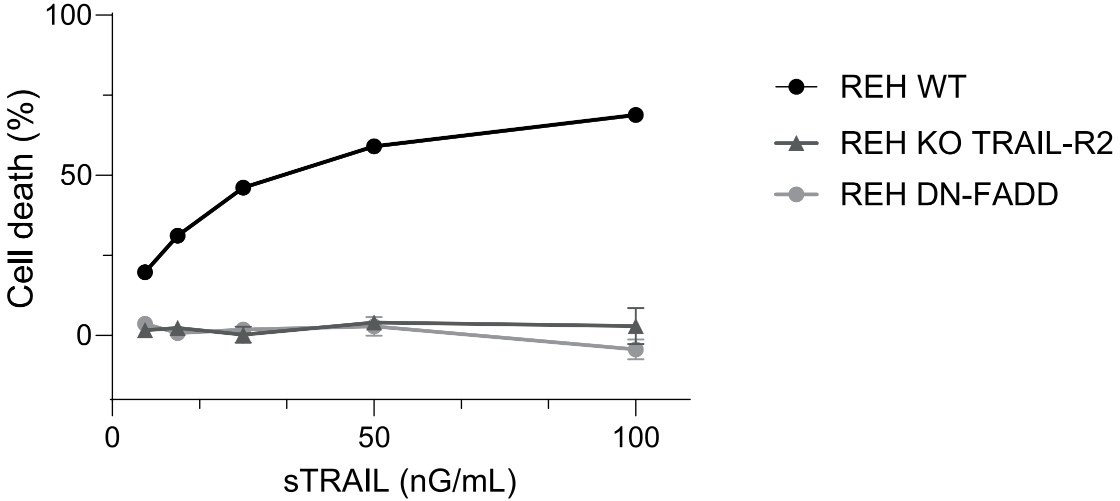


**Supplemental Figure S1: TRAIL-R2 KO and DN-FADD expression impair the apoptosis induced by sTRAIL**
Indicated cell lines were cultured in the presence of increasing concentrations of soluble TRAIL (sTRAIL) for 24h. Cell death was measured by flow cytometry after PI staining. Means of 3 independent experiments are presented with SEM.
