## Supplemental methods for "TRAIL orchestrates ThINKK-induced NK cell cytotoxicity against childhood acute lymphoblastic leukemia"

**697 transduction with *TNFRSF10B* expressing lentiviral vector**

Plasmids

The pcDNA3-TRAIL-R2 plasmid was a gift from Francis Chan (Addgene plasmid # 61383 ; http://n2t.net/addgene:61383 ; RRID:Addgene_61383)^1^. the pMD2.G (VSV-G) plasmid (Addgene plasmid#12259; http://n2t.net/addgene:12259; RRID:Addgene_12259) as well as the psPAX2 plasmid (Addgene plasmid # 12260 ; http://n2t.net/addgene:12260 ; RRID:Addgene_12260) were a gift from Didier Trono and the pHRSIN-UCOE0. 7-SFFV- DEST-PGK-BSD plasmid and the pENTR1A plasmid (Invitrogen, 11813-011) were a gift from Elie Haddad.

Construction of the gateway entry vector and lentivirus vector

A unique NotI restriction site was added at the 3’ end of the TRAIL-R2 cDNA on the pcDNA3 plasmid by PCR with primers: TRAILR2-pcDNA3-Fwd (TAATCTGGATCCGCCATGGAAC) and TRAILR2-pcDNA3-Rev (ACTAATGCGGCCGCTTAGGACATGGCAGAGTCT) (both from IDT), using Platinum SuperFi PCR Master Mix (Life Technologies, 12358-002) for 35 cycles and a melting temperature of 60°C on a Veriti 96 well thermal cycler. The PCR fragment (TRAIL-R2 cDNA) was digested using the BamHI-HF (NEB, # R3136S) and the NotI-HF (NEB, #R3189S) enzymes, as was the pENTR1A plasmid.

The digested PCR fragment and plasmid were loaded onto a Low Melting Agarose (Life Technologies, 16520-050) gel and purified with the MinElute Gel Extraction Kit (Qiagen, 28604). The digested PCR fragment was then ligated to the digested plasmid using T4 Ligase (NEB, #M0202S) at a 3:1 ratio (90 fmol of fragment for 30 fmol of plasmid). 3 µL of the ligation reaction was then added to 50 µL of TOP10 competent cells (Life Technologies, C4040-03) and spread on LB [Lennox] agar plates with Kanamycin, 25µg/ml (Life technologies, 15160-054). After overnight incubation at 37°C, a few colonies were chosen and plasmid DNA isolated using QIAprep Spin Miniprep kit (QIAGEN, 27104). This plasmid DNA was then sent to the IRIC's Genomics Platform for sequencing.

Finally, one of the plasmid DNAs with the correct TRAIL-R2 cDNA sequence was used to transfer the cDNA from the pENTR1A plasmid (100ng) to the pHRSIN-UCOE0.7-SFFV- DEST-PGK-BSD plasmid (150 ng) using the Gateway LR Clonase II Enzyme Mix Kit (ThermoFisher, 11791-020) per manufacturer’s instructions. One tenth of this reaction was then added to 50 µL of Stabl3 competent cells (Life Technologies, C737303) and spread on LB [Lennox] agar plates with Ampicillin, 100µg/ml (Wisent, 450110-XL). After overnight incubation at 37°C, a few colonies were chosen and plasmid DNA isolated using the QIAprep Spin Miniprep kit (QIAGEN, 27104). Plasmid size was verified on an agarose gel, after digestion with BamHI-HF (NEB, # R3136S). A larger quantity of the plasmid DNA was prepared using the HiSpeed Plasmid Maxi Kit (QIAGEN, 12662).

Lentivirus production

The TRAIL-R2 lentiviral particles (LVs) were generated by the transient cotransfection of HEK 293T/17 cells with a three-plasmid system. HEK 293T/17 cells (CRL-11268, ATCC), obtained from ATCC (Manassas, VA, USA), were plated in 10cm plates (2.6x10^6^ cells) and covered with 8ml of DMEM (Wisent Inc.) + 10% FBS (Wisent Inc) without antibiotics. The next day, the lentiviral transfer vector plasmid (pHRSIN-UCOE0.7-SFFV-TRAILR2-PGK-BSD ; 8.6 μg), packaging plasmid (psPAX2; 8.6 μg) and envelope plasmid (pMD2.G; 7 μg) DNA were mixed with Opti-MEM media (Invitrogen Life Technologies). PEI (Polyethylenimine MW 25000, Linear (Polysciences, 23966-100)) (24.2µg) was diluted with Opti-MEM to a volume of 1ml, using the formula previously described by Girard-Gagnepain et al^2^. PEI was added slowly to the DNA mixture from above and incubated at room temperature for 20 min. The DNA/PEI mixture was then dropped slowly on the HEK cells incubated at 37°C in a 5% CO_2_ humidified atmosphere for 24 hours. The media was then replaced with fresh Opti-MEM and incubated another 48 hours (72h post-transfection). The supernatants containing LVs were collected, and concentrated with PEG-it Virus Precipitation Solution (System Biosciences, LV810A-1) according to manufacturer’s instructions. Titration of the lentivirus was performed using the HIV-1 p24 ELISA Assay (XpressBio, XB-1000) according to manufacturer’s instructions.

Generation of *TRAIL-R2* (*TNFRSF10B*) knock-in cell line

697 cells were plated in RPMI 1640 medium (Wisent Inc.) supplemented with 10% FBS (Wisent Inc.). TRAIL-R2 lentiviral particles were added at different MOIs (1, 5, 10 and 20) in medium containing Protamine Sulphate (Fresenius Kabi, C22905). The cells were then incubated at 37°C in a 5% CO_2_ atmosphere. Multiple washes were performed 72 hours after transduction, and cells were resuspended in culture medium. Samples of each culture were analysed by flow cytometry using a PE anti-TRAIL-R2 antibody (Biolegend).

TRAIL-R2 expressing 697 cells were then sorted using a clonal selection process to establish a single-cell based colony. Twelve clones were then characterized in respect to their expression of TRAIL-R2 and the one with the highest MFI was selected and use for all NK cell functional assays. The genomic sequencing of six clones was performed to confirm the integrity of the transgene in these clones. Surprisingly, all the clones exhibit mutations or deletion in the cytoplasmic death domain of the protein, making these receptors unable to induce cell death upon their interaction with their ligand TRAIL. We decided to use the E10-697 clone that carries a stop codon at position 234 and therefore lacks the entire cytoplasmic region. Our Western blot analysis confirmed the truncated form of TRAIL-R2 (Supplemental Figure S1) while flow cytometry analysis confirmed the presence of TRAIL-R2 at the surface membrane (Figure 5A).

**TNFRSF10B knock-out using CRISPR-Cas9 technology**

Plasmids

LentiCas9-Blast (Addgene plasmid # 52962 ; http://n2t.net/addgene:52962 ; RRID:Addgene_52962) and lentiGuide-Puro (Addgene plasmid # 52963 ; http://n2t.net/addgene:52963 ; RRID:Addgene_52963) plasmids were provided by the Gene Editing Platform of the Azrieli CHU Sainte-Justine Research Center and a gift from Feng Zhang^3^. LentiCas9-Blast and pLentiGuide are plasmids with a lentiviral backbone that express human codon-optimized Streptococcus pyogenes Cas9 protein along with the blasticidin resistance gene and CRISPR chimeric RNA element with the puromycin resistance gene, respectively.

Generation of *TNFRSF10B* knock-out REH cell line

The Gene Platform of the Azrieli CHU Sainte-Justine Research Center developed the REH KO TRAIL-R2 cell line. Briefly, REH cells were transduced with with the LentiCas9-Blast LV (lentivirus). Blasticidin (5 ng/ml) (Wisent Inc.) was added to culture media following transduction to select for caspase-9 (Cas9)-expressing REH cells (REH-Cas9-WT). The production of Cas9 protein by wild-type REH cells (REH-Cas9-WT) was confirmed by western blotting.

REH-Cas9-WT cells were then transduced with the pLentiGuide LV expressing 3 different gRNAs. Puromycin (1 μg/ml) (Wisent Inc) was used to select for the cells expressing the TNFRSF10B sgRNAs. A mismatch assay with the GeneArt Genomic Cleavage Detection Kit (Life Technologies) was performed and all 3 gRNAs were found to have a 15% efficiency. The culture with gRNA #3 was used for further clonal selection.

Flow cytometry using a PE anti-human TRAIL-R2 antibody (Biolegend) was performed on the cell culture and only cells negative for TRAIL-R2 expression (REH-Casp9-TRAIL-R2) relative to the REH-Cas9-WT cells were retained. Clonal selection and expansion was performed by sorting single cells lacking TRAIL-R2 expression (REH KO TRAIL-R2) on a BD FACSAria II sorter.
